# Pathway specific drive of cerebellar Golgi cells reveals integrative rules of cortical inhibition

**DOI:** 10.1101/356378

**Authors:** Sawako Tabuchi, Jesse I. Gilmer, Karen Purba, Abigail L. Person

**Affiliations:** Department of Physiology & Biophysics; Neuroscience Graduate Program, University of Colorado Denver University of Colorado School of Medicine Aurora, CO 80045

## Abstract

Cerebellar granule cells (GrCs) constitute over half of all neurons in the vertebrate brain and are proposed to decorrelate convergent mossy fiber inputs in service of learning. Interneurons within the granule cell layer, Golgi cells (GoCs), are the primary inhibitors of this vast population and therefore play a major role in influencing the computations performed within the layer. Despite this central function for GoCs, few studies have directly examined how GoCs integrate inputs from specific afferents which vary in density to regulate GrC population activity. We used a variety of methods in mice of either sex to study feedforward inhibition recruited by identified MFs, focusing on features that would influence integration by GrCs. Comprehensive 3D reconstruction and quantification of GoC axonal boutons revealed tightly clustered boutons that focus feedforward inhibition in the neighborhood of GoC somata. Acute whole cell patch clamp recordings from GrCs in brain slices showed that despite high bouton density, fast phasic inhibition was very sparse relative to slow spillover mediated inhibition. Furthermore, dynamic clamp simulating inhibition combined with optogenetic mossy fiber activation supported the predominant role of slow spillover mediated inhibition. Whole cell recordings from GoCs revealed a role for the density of active MFs in preferentially driving them. Thus, our data provide empirical conformation of predicted rules by which MFs activate GoCs to regulate GrC activity levels.

## Significance Statement

A unifying framework in neural circuit analysis is identifying circuit motifs that subserve common computations. Widefield interneurons are a type of inhibitory interneuron that globally inhibit neighbors that have been studied extensively in the insect olfactory system and proposed to serve pattern separation functions. Cerebellar Golgi cells (GoCs), a type of mammalian widefield inhibitory interneuron observed in the granule cell layer, are well suited to perform normalization or pattern separation functions but the relationship between spatial characteristics of input patterns to GoC mediated inhibition has received limited attention. This study provides unprecedented quantitative structural details of GoCs and identifies a role for population input activity levels in recruiting inhibition using in vitro electrophysiology and optogenetics.

## Introduction

A fundamental function of the cerebellar granule cell (GrC) is to decorrelate information conveyed via convergent multimodal mossy fibers (MFs), increasing utility for learned associations (Marr, 1969; Albus, 1971; Billings et al., 2014; Cayco-Gajic et al., 2017). Recent work has demonstrated that GrCs receive and respond to MFs conveying diverse information (Huang et al., 2013; Ishikawa et al., 2015) but little attention has been paid to the potential role of multimodal integration by Golgi cells (GoCs). GoCs are in a key position to regulate expansion recoding by GrCs since feedforward inhibition sets spiking threshold and thereby the number of different afferents required to drive GrC firing (Marr, 1969; D’Angelo et al., 2013). Indeed, theory suggests that feedforward inhibition via GoCs performs a thresholding-like function, clamping the number of active GrCs at a relatively fixed level by engaging GoCs in a scaled manner with increasing activity from MFs (Marr, 1969; Medina et al., 2000)

GoC inhibition of GrCs has been studied extensively in slices, and is characteristically diverse. Fast phasic IPSCs, a pronounced slow spillover-mediated component, and ‘tonic’ GABA_A_-receptor mediated currents are all forms of inhibition mediated by GoCs (for review, see Farrant and Nusser, 2005). The spill-over and tonic inhibitory tone within the layer would seemingly provide an ideal mechanism for widely inhibiting the vast number of GrCs without necessarily forming direct contact with each cell. Nevertheless, fast phasic feedforward IPSCs have been observed with electrical stimulation paradigms and suggested to sharpen GrC responses to MF inputs, producing a temporal-windowing effect analogous to feedforward temporal sharpening of thalamic input in neocortical pyramidal neurons (Pouille and Scanziani, 2001; Farrant and Nusser, 2005; D’Angelo and De Zeeuw, 2009; Nieus et al., 2014). This diversity of potential computations mediated by a single cell, raised the question of whether these particular features of GoC transmission are selectively engaged by specific afferent streams or whether they are part of a continuum of inhibitory transmission onto GrCs that result from probabilistic innervation of GrCs within glomeruli. Furthermore, relating GoC recruitment to the density of active MFs is critical for testing the hypothesis of dynamic thresholding in service of pattern separation.

Another challenge for GoCs is inhibiting the vast number of GrCs to regulate activity within the granule cell layer. GoC axons are famously dense, but details of spatial ramification patterns that define the likelihood of local GrCs sharing inhibition remain undefined. Indeed, the problem of quantitatively addressing the distribution of inhibition from a single Golgi cell was described by Ramon y Cajal: “When one of these axons appears completely impregnated in a Golgi preparation, it is almost impossible to follow its complete arborization.… It is only in the incomplete impregnations of adult animals … that one can study the course and divisions of the axon. Ramon y Cajal 1890a” (Palay and Chan-Palay, 1974). To our knowledge, this observation remains relevant in contemporary literature where all GoC reconstructions have been incomplete (Simpson et al., 2005; Barmack and Yakhnitsa, 2008; Kanichay and Silver, 2008; Vervaeke et al., 2010; Vervaeke et al., 2012; Szoboszlay et al., 2016; Valera et al., 2016).

To address these questions, we used a variety of methods to resolve GoC connectivity rules and the capacity of specific afferents to produce fast phasic and slow spill-over mediated inhibition. We performed comprehensive single cell, high-resolution reconstruction of GoCs with quantitative morphological analysis to estimate glomerular innervation by GoCs. Optogenetic activation of specific MF afferents from the pontine or cerebellar nuclei, which differ systematically in their density, were used with electrophysiological recordings of GoCs from slices to test the prediction that the density of afferent activity selectively engages inhibitory mechanisms to regulate GrC threshold. Combining MF optogenetic activation with dynamic clamp mimicking feedforward fast or slow phasic inhibition, supported the view that spillover-mediated inhibition selectively modulates GrC activity and challenges predictions of temporal windowing in GrCs by phasic FFI.

## Materials and Methods

### Animals

All procedures followed the National Institutes of Health Guidelines and were approved by the Institutional Animal Care and Use Committee and Institutional Biosafety Committee at the University of Colorado Anschutz Medical Campus. Animals were housed in an environmentally controlled room, kept on a 12 h light/dark cycle and had *ad libitum* access to food and water. Adult mice of either sex were used in all experiments; sex was not monitored for experimental groupings. Genotypes used were C57BL/6 (Charles River Laboratories), Neurotensin receptor1-Cre (Ntsr1-Cre; Mutant Mouse Regional Resource Center, STOCK Tg (Ntsr1-cre) GN220Gsat/Mmucd) and GlyT2-eGFP (Salk Institute; Zwilhofer et al., 2005; Tg(Slc6a5-EGFP)13Uze). All transgenic animals were bred on a C57BL/6 background and maintained as heterozygotes. Ntsr1-Cre animals were genotyped for Cre, and GlyT2-eGFP animals were genotyped for eGFP (Transnetyx).

### Virus Injections

For surgical procedures, at least one month old mice were anesthetized with i.p. injections of ketamine hydrochloride (100 mg/kg) and xylazine (10 mg/kg) cocktail. Mice were placed in a stereotaxic apparatus and bupivicaine (6 mg/kg) was injected along incision line. Craniotomies were made above the cerebellar cortex (from lambda: −1.9 mm, 1.1 mm lateral, 1.2 mm ventral), interposed nuclei (IN) (from lambda: −1.9 mm, 1.1 mm lateral, 2.4 mm ventral) and pontine nuclei (from bregma: −5.4 mm, 0.5 mm lateral, 5.2 mm ventral). Pressure injections of virus were administrated using a pulled glass pipette (7-9 μm tip diameter). Mice were allowed to survive for more than 6 weeks before experiments, which we found in pilot experiments optimized expression of reporter proteins in MF terminals.

### Viruses

AAV8-hSyn1-mCherry-Cre (titer: 10^2^, UNC) and AAV2-CAG-FLEX-eGFP (titer: 10^12^, Penn) were coinjected to cerebellar cortex to sparsely label neurons for morphological analysis of Golgi cells. AAV2-hSyn1-hChR2(H134R)-mCherry-WPRE (University of North Carolina Vector Core) were injected to wildtype mouse IN and pontine nuclei to induce ChR2 expression for electrophysiological recordings. AAV2-EF1a-DIO-hChR2(H134R)-mCherry-WPRE-pA was injected into the IN of Ntsr1-Cre mice for a subset of nucleocortical MF studies (University of North Carolina Vector Core).

### Electrophysiology

#### Slice preparation

At least 6 weeks after virus injection, mice were deeply anesthetized with isoflurane before transcardial perfusion with warm (37-40 °C), oxygenated (95% O_2_-5% CO_2_) Tyrode’s solution containing (in mM): 123.75 NaCl, 3.5 KCl, 26 NaHCO_3_, 1.25 NH_2_PO_4_, 1.5 CaCl_2_, 1 MgCl_2_ and 10 glucose. Dissected cerebellum was sliced in 300 μm in coronal plane for Golgi cell recordings and either parasagittal or coronal plane for granule cell recordings on a Leica VT1000S Vibratome. Slices were transferred to an oxygenated Tyrode’s solution (37 °C) and incubated for 30-60 min.

#### In vitro recordings

One hour after slicing for granule cell recordings and immediately after slicing for Golgi cell recordings (Hull and Regehr, 2012), tissue was transferred to the recording chamber. Oxygenated Tyrode’s solution (30 °C) was perfused over the slice at 3 ml/min and visualized with Zeiss AxioExaminer equipped with xenon lamp LAMBDA DG-4 (Sutter Instrument) for optogenetic stimulation through the objective. MFs were stimulated optogenetically with 2 ms light pulses with a measured power of 18.5 mW (3.77 mW/mm^2^) at 473 nm. Pulled glass patch electrodes (Golgi cell: 2-3.5 ΜΩ, Granule cell: 4 - 6 ΜΩ; Sutter Instruments, P-97) were filled with K-gluconate-based internal solution contained the following (in mM): 132 K-gluconate, Na-gluconate, 4.4 NaCl, 2.2 MgCl_2_, 1.1 EGTA, 11 EGTA, 14 TRIS creatine phosphate, 4 Mg-ATP and 0.3 Tris-GTP. For all Golgi cell recordings and a subset of granule cell recordings, biocytin (0.3%; Tocris Bioscience) was added to the internal solution. Whole-cell recordings were made in current-clamp and voltage-clamp mode, low-pass filtered at 6-10 kHz, amplified with a MultiClamp 700C, partially compensated for series resistance, digitized at 10-50 kHz with a Digidata 1550 and monitored with pClamp acquisition software (Molecular Devices). Blockade of neurotransmitter receptors was achieved with bath application of 10 μM RS-CPP (Tocris Bioscience) to block NMDAR-mediated currents, and 10 μM SR95531 (Tocris Bioscience) to block GABA_A_R-mediated currents. Data were analyzed with custom routines and with the Neuromatic package (ThinkRandom) in IGOR Pro (Wavemetrics; RRID: SCR_000325).

#### Dynamic clamp

We built and utilized a custom microcontroller-based dynamic clamp system with 10 μs input-output latency (Desai et al., 2017). We modified the layout, customized resistors, and made a custom 3D printed enclosure, and calibrated the system with details provided as an open resource from the Optogenetics and Neural Engineering Core at the University of Colorado Anschutz Medical Campus (dynamicclamp.com). IPSC parameters were calculated from our GrCs recording data set or previous reports (Rossi and Hamann, 1998). Fast phasic IPSCs had a 2.15 ms rise time, 2.29 ms decay tau, and a peak conductance of 0.3 nS or 1.2 nS. Slow IPSCs: had a 30.2 ms rise time, 630 ms decay tau, and peak conductances of 0.032 or 0.32 nS. IPSG timing was controlled by a Master-8 Pulse Stimulator (A.M.P.I).

### Tissue preparation for light microscopy

Mice were overdosed with an intraperitoneal injection of a sodium pentobarbital solution, Fatal Plus (Vortech Pharmaceuticals), and perfused transcardially with 0.9% saline followed by 4% paraformaldehyde. Brains were removed and postfixed for at least 24 hrs then cryoprotected in 30% sucrose. Tissue was sliced in 40 μm serial coronal sections using a freezing microtome and stored in 0.1 M PB.

For post hoc morphological analyses following slice electrophysiology, tissue was transferred to warm 4% paraformaldehyde and postfixed for less than 3 hrs then placed in 30% sucrose. Tissue was rinsed in 0.1M PB for 30 min then treated in 0.3% Triton X-100 in 0.1M PB for 2 hrs followed by three washes in PB (10 min each). To visualize biocytin, tissue was incubated with streptavidin conjugated to AlexaFluor 647 or 555 (Life Technologies) diluted 1:100 in 0.1 M PB overnight at 4 °C followed by three washes in PB (20 min each).

### Imaging

Confocal images were obtained using Zeiss LSM 780. To reconstruct all processes and/or map all axonal boutons of GoCs, we imaged in fine z stacks (0.39 μm stacks) through Plan-Apochrom 63x/1.4 oil DIC M27 objective by Ar-Iron laser. For MF density analysis images of filled cells were collected and density analyzed within 53615±5180 μm^2^ of the GoC soma. Images examining MF proximity to GoCs were visualized in Zen software using transparent rendering mode. High resolution imaging for GlyT2-GFP boutons was performed with a Nikon A1r-HD confocal with a Plan Apo 60x oil objective, NA 1.4. Deconvolution was performed in NIS-Elements C imaging software.

### Morphological analysis

GoC reconstructions were performed using Neurolucida 360 software (MBF Bioscience; RRID: SCR_001775). Processes were traced in user-guided mode. Fine grained reconstructions captured axonal and dendritic swellings by adjusting points along processes to match thickness. Because individual GoCs spanned multiple sections, each section was individually reconstructed, then each reconstructed section was stacked using morphological landmarks visible across sections, such as the Purkinje cell layer. To define the relative extent of basal axons and dendrites, we traced these processes with attention to process thickness and contours. Axons were characterized by their smaller diameter (~0.2 μm) and were studded with boutons (~3 μm diameter). For all analyses of reconstructed GoC morphological features, we used Neurolucida Explorer. To define the GCL volume occupied by the cell, we computed the convex hull volume which calculates the volume of a convex polygon connecting the tips of the distal processes. To map boutons, we used Neurolucida 360 and placed markers on each bouton by with 0.39 μm z resolution. All images were aligned and coordinates of boutons were exported and processed in Matlab (RRID: SCR_001622). Nearest Boutons: All distance analyses used Euclidean distance. Nearest bouton color maps were constructed as 3D-scatter plots in Matlab. Bouton distance probabilities were computed in 0.2 μm bins and normalized by the total number of boutons. Comparisons of bouton density between single Golgi cells and global GlyT2-GFP label was calculated by first computing the local density of boutons for single GoCs, defined as the number of boutons within a 25 μm radius sphere of each bouton. The density of GlyT2 boutons was determined by mapping boutons in 6 GlyT2-GFP GCL samples and dividing the total bouton count by the volume of the image. Density measurements were then compared between single GoCs and the GlyT2-GFP+ population by computing the ratio of bouton densities in a single GoC and mean GlyT2-GFP bouton density. The ratio was used to estimate the fraction of bouton density accounted for by a single GoC. We display the distribution of density ratios in a histogram normalized by the total number of boutons with a bin size of 0.05.

### Experimental design and Statistical analysis

Paired and unpaired t-test were performed using RStudio (version 1.0.136; RStudio). Linear regression analysis was performed using GraphPad Prism 7.04 (GraphPad software; RRID:SCR_002798). Statistical tests are specified in the text. Electrophysiological and morphological analyses are described above.

## Results

### Spatial distribution of Golgi cell processes

Rules of integration in GrCs will depend critically on the structure of feedforward inhibition within the granule cell layer (GCL). Previous studies have proposed contradictory integrative models. Influential older work proposed that GoC axons tile the GCL in a non-overlapping manner such that GrCs are innervated by a single GoC (Eccles et al., 1967), yet physiological measurements estimated convergence of multiple GoCs onto GrCs but the existence of slow spillover current makes convergence estimates challenging (Rossi and Hamann, 1998). Several morphological features of GoCs, including axonal volume and bouton density, are each essential to infer the inhibitory convergence in the GCL. Therefore, we set out to define the density of GoC basal axons to test these assumptions and inform inhibitory connectivity rules in the GCL. We performed comprehensive basal arbor reconstruction to quantify GoC morphological characteristics that contribute to spatial distribution of feedforward inhibition in the GCL. To do so, we used a sparse viral labeling technique which restricted the number of fluorescent GoCs in cerebellar cortex. A combination of low titer Cre recombinase-expressing virus (AAV8-hSyn1-mCherry-Cre; titer: 10^2^) and high titer Cre-dependent reporter virus (AAV2-CAG-FLEX-eGFP; titer: 10^12^) were injected into cerebellar cortex which resulted in very sparse labeling of individual GoCs (Fig. 1A).

**Figure 1.**
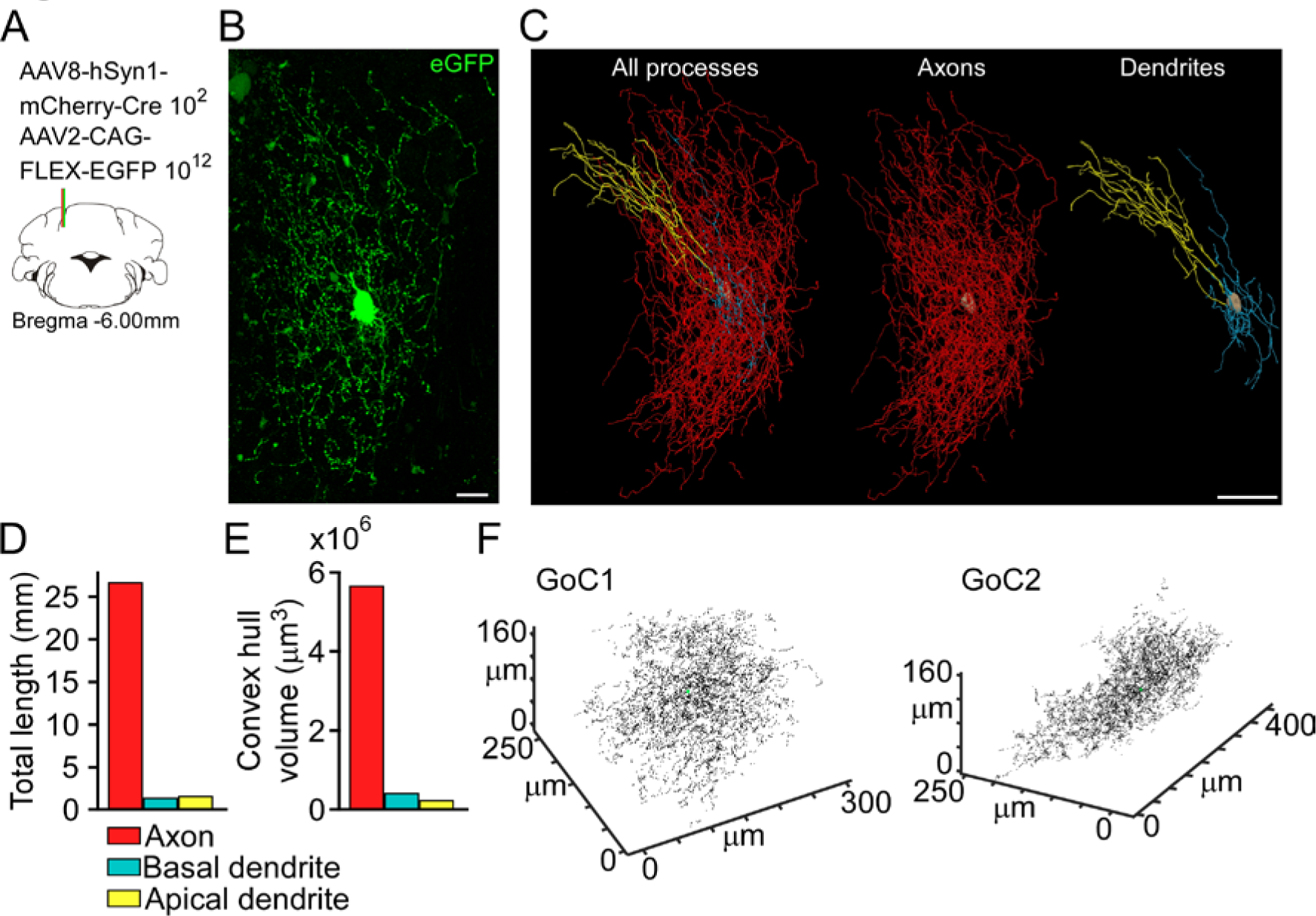
Sparse viral label and comprehensive reconstruction of cerebellar GoCs. **A**, Schematic of sparse viral labeling technique where low titre AAV8-hSyn1-mCherry-Cre 10 and high titre AAV2-CAG-FLEX-EGFP 10^12^ were coinjected to cerebellar cortex. **B**, Example of sparsely labeled GoC. Scale: 20 μm. **C**, Comprehensive, 3D reconstruction of GoC displayed in 2D. Red, axon; Cyan: basal dendrite; Yellow: apical dendrite, Beige, soma. Left: overview of the single GoC. Middle: axon processes in GCL. Right: apical dendrites and basal dendrites. Scale: 50 μm. **D**, Total length of each process type. **E**, Convex hull volume of each process type. **F**, Maps of GoC axon boutons of sparsely labeled GoCs. Black dots indicate the location of each bouton and green circle indicates soma.

In keeping with previous reports, sparse GoC label revealed extensive basal axons, which were characterized by their small diameter (~0.2 μm) (Palay and Chan-Palay, 1976; Holtzman et al., 2006; Barmack and Yakhnitsa, 2008; Hull and Regehr, 2012; Vervaeke et al., 2012; Ankri et al., 2015; Valera et al., 2015). An example GoC is shown in Fig. 1B. This GoC extended processes ~200 μm mediolaterally, ~330 μm dorsoventrally, and ~180 μm rostrocaudally, occupying a volume of approximately 5.7×10^6^ μm^3^ and overall axonal length of 26.7 mm (Neurolucida; see Methods). This volume was dominated by axonal processes, which defined the boundaries of this field. Therefore, this GoC basal axon encompasses a volume that contains approximately 15,000 GrCs and 580 MF rosettes (MFRs) based on estimates of GrC and MFR densities (Palkovits et al., 1971). GoC basal dendrites were distinguishable from axons by their larger diameter (0.3 - 3.2 μm) and smooth surface devoid of boutons as previous described (Palay and Chan-Palay, 1974; Hull and Regehr, 2012; Vervaeke et al., 2012; Ankri et al., 2015; Rudolph et al., 2015; Szoboszlay et al., 2016). Dendrites comingled with axons but were considerably shorter (Fig. 1C, right), with a length totaling 1.4 mm (Fig. 1D) and encompassing a computed volume of 0.41 × 10^6^ μm^3^ (Fig. 1 E; convex hull volume; Neurolucida, see Methods). Because the total axon processes was ~20 times longer than basal dendrites, GoC mediated inhibition is predicted to be distributed more widely than the afferent input in the GCL, as has been previously noted (D’Angelo et al., 2013).

The highly restricted cell labeling *in vivo* also permitted novel quantification of GoC bouton density and numbers. Axons also possessed distinct bouton swellings (~3 μm diameter). We comprehensively mapped boutons of two GoCs located in vermal lobule 4/5 and quantified bouton density on an additional partially reconstructed GoC (Fig. 1F). Axons of individual GoCs were studded with a total of 6,000-7,500 boutons (5.9×10^3^ and 7.46×10^3^), for an average single GoC bouton density of 3.25×10^5^ boutons/mm^3^. We calculated the nearest bouton by Euclidean distance, which revealed that a majority (84 - 93%) were within 4 μm of one another with the relatively uniform distribution of nearest boutons illustrated in the color map of nearest neighbors (Fig. 2A-C; median nearest bouton, 2.18-2.80 μm; n=13,360 boutons). Although the nearest boutons remained fairly constant across the axonal arbor, visual inspection of the bouton maps suggested clustering toward the soma. We therefore analyzed the number of neighbors within 10 μm of each bouton. This analysis revealed highly structured clustering, where a majority of boutons close to soma had more than 20 close neighbors, with density dropping sharply as a function of distance from the soma (Fig. 2D, E).

**Figure 2.**
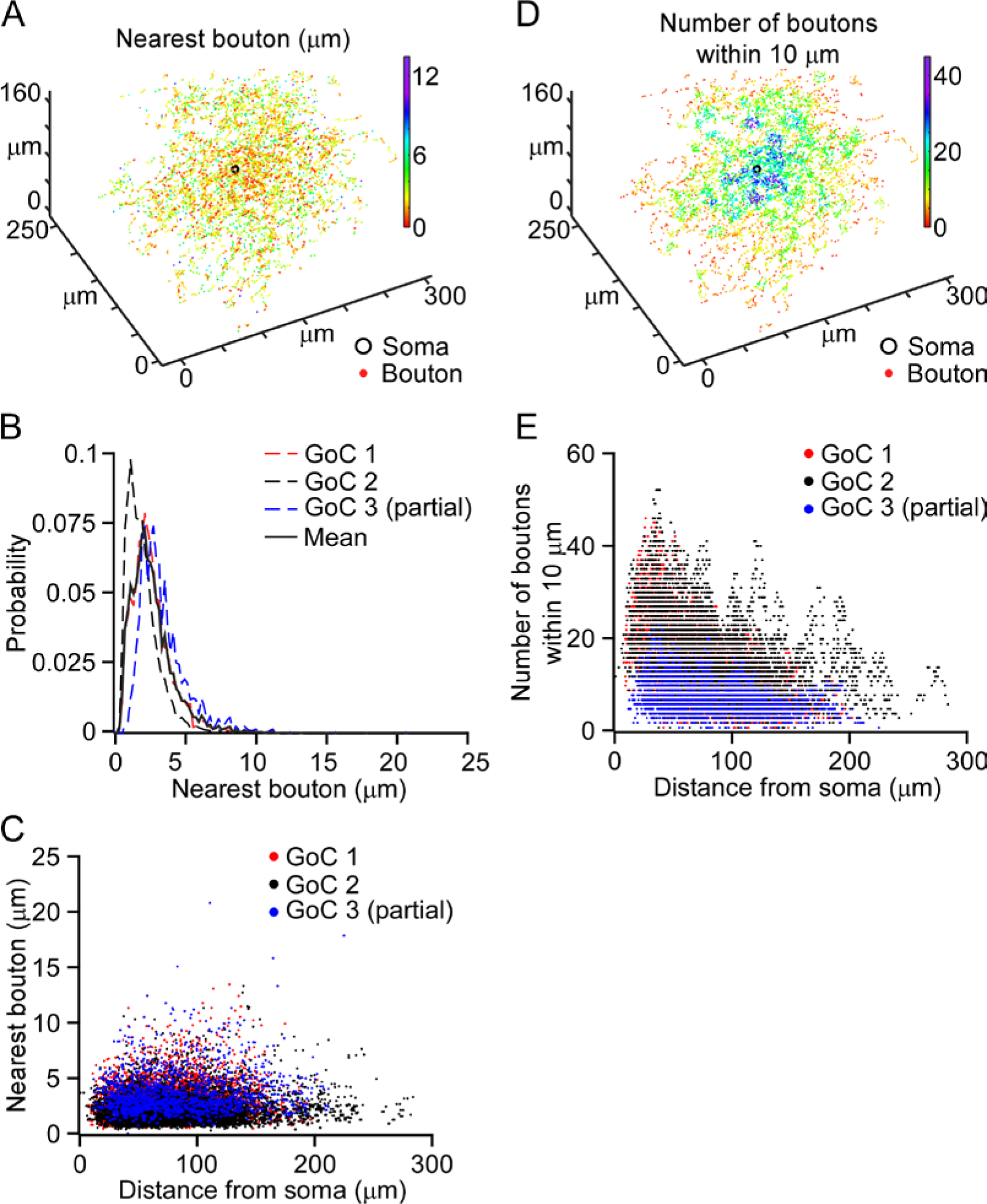
Clustered boutons revealed with high resolution mapping. **A**, Bouton map of GoC with distance to the nearest bouton represented by color. **B**, Probability histogram of nearest bouton distances in three GoCs (dashed) and mean of three cells (solid). Bin width: 0.2 μm. **C**, Summary of nearest boutons as a function of distance from soma. **D**, Bouton map of the same GoC shown in *>A*> with the number of boutons within 10 μm radius from each boutons represented by color. **E**, Number of boutons within 10 μm radius from each boutons as a function of distance from the soma.

These comprehensive bouton maps allowed us to estimate GoC overlap at the glomerulus. To do so, we first compared bouton distributions from single cell reconstructions with those observed in GlyT2-eGFP mice, which label approximately 85% of GoCs (Fig. 3A) (Simat et al., 2007). Mapping boutons in these preparations revealed an overall bouton density of 6.54×10^6^ boutons/mm^3^. Distributions of nearest boutons differed slightly between single cell reconstructions and GlyT2-eGFP label (Fig. 3B). Not surprisingly, GlyT2-eGFP labeled boutons were more closely spaced, with a prominent peak at ~ 1.5 μm compared to the peak at 3 μm for the single neurons. Differences between single cells and the population were more pronounced when looking at local clustering, seen by measuring the distances to the closest 10 neighbors (Fig. 3C). GlyT2-eGFP labeled boutons were denser, with the closest 10 boutons appearing on average ~4 μm away from every other bouton. By contrast, the nearest 10 boutons to each bouton on axons from individual neurons averaged ~7 μm. To estimate GoC axonal overlap, we next examined the proportion of GlyT2 bouton density accounted for by a single GoC. In areas within 50 μm of a GoC soma, single GoC axons tended to account for 60-80% of bouton density, suggesting that glomeruli here are likely dominated by a single GoC (Fig. 3D). The proportion of population bouton density accounted for by a single GoC falls off with distance from the GoC soma (Fig. 3D). Despite this focal clustering of GoC boutons near the soma, based on the density of GlyT2 labeled boutons accounted for by a single GoC, a plurality of GoCs boutons mingle with boutons from other GoCs, such that at any given site, approximate 20% of boutons are from a single GoC, suggesting that typically 5 GoCs overlap (Fig. 3E).

**Figure 3.**
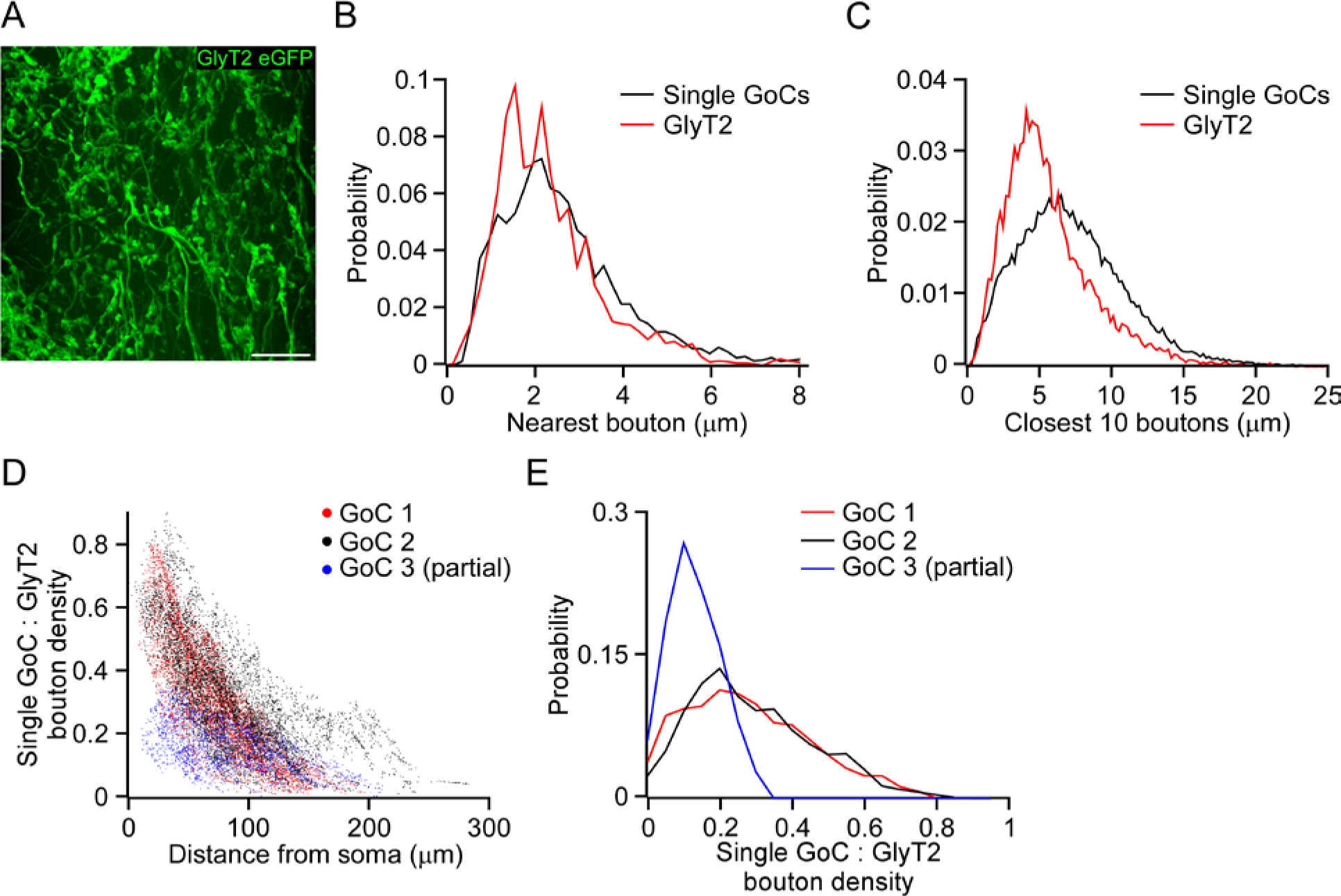
Contribution of single GoCs to population distribution. **A**, GlyT2-eGFP expressing processes in mouse GCL. Scale: 10 μm. **B**, Probability histograms of nearest bouton distances for single GoCs (black) and the population of GoCs labeled with GlyT2-GFP (red). Bin width: 0.2 μm. **C**, Probability histograms of average distances to the closest 10 boutons from each bouton for single GoCs (black) and a population of GoCs labeled with GlyT2-GFP (red). Bin width: 0.2 μm. **D**, Ratio of single GoC: population bouton density as a function of distance from GoC soma. The mean GlyT2 bouton density within a 25 μm radius sphere was compared to the number of boutons within a 25 μm radius sphere for each bouton in the single GoCs. **E**, Probability histogram of density ratio for single GoC: population for each bouton, indicating that boutons from a single GoC typically account for less than half of the local bouton population. Bin width: 0.05.

### Low probability fast phasic feedforward inhibition in the GCL

The spatial extent of GoC processes position them as sites of MF convergence and inhibitory divergence, motivating experiments examining how feedforward inhibition is recruited by identified cerebellar inputs. To begin to test how diverse MF afferents interact in the GCL, we first scaled down the question to address how identified sources of information recruit GrCs and feedforward inhibition via GoCs. We examined uni-modal information processing of GrCs by recording postsynaptic currents evoked by optogenetic stimulation of a subset of MFs originating from neurons in the cerebellar nuclei, known as the nucleocortical (NC) MFs or those originating in the pontine nuclei. AAV2-hSyn1-hChR2-mCherry-WPRE-PA (See Methods) was injected into either the interposed nucleus (IN) or pontine nuclei to express ChR2 in MFs. We examined the GrC responses to MF optogenetic stimulation in an acute brain slice preparation from adult mice. We isolated EPSCs and IPSCs by holding GrCs at −70 mV or 0 mV respectively and stimulated ChR2 expressing MFs at between 20 - 60 Hz with blue light pulses delivered through the objective (Fig. 4A; 2 ms pulses; 0.96 mW/mm^2^).

**Figure 4.**
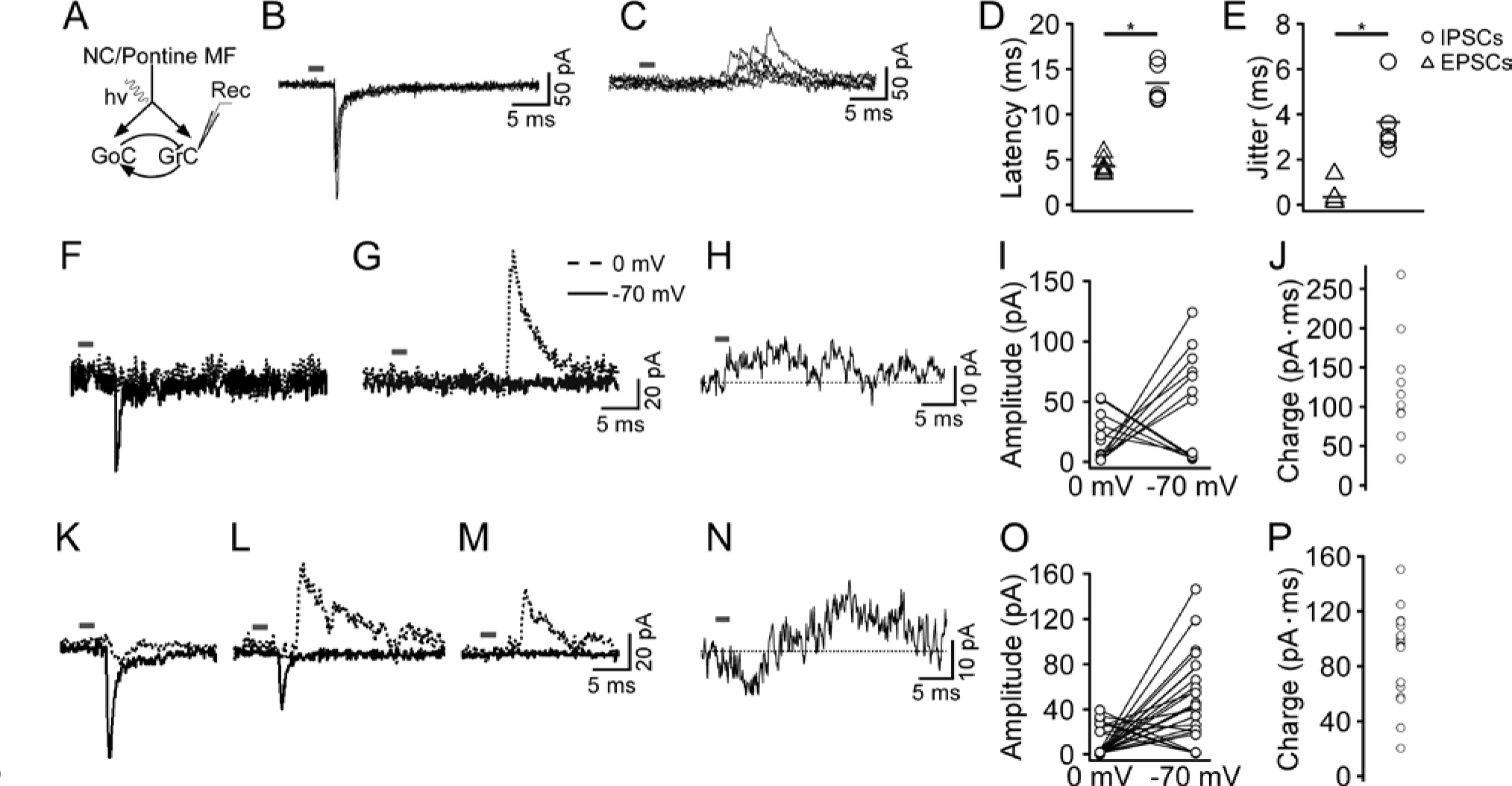
GrC responses to optogenetic activation of identified MF populations. **A**, Schematic diagram of recording configuration. ChR2-expressing NC or pontine MFs were optogenetically stimulated during GrC recordings. **B and C**, Overlaid EPSCs or IPSCs evoked by 2 ms optogenetic stimulation of NC MFs at −70 or 0 mV holding. **D**, Latency of EPSCs and fast phasic IPSCs (p < 0.001; unpaired t-test, gray bar = mean). **E**, Jitter of timing of EPSCs and fast phasic IPSCs was significantly different (p < 0.001, unpaired t-test; bar = mean). **F and G**, Representative traces of responses to light stimulation following NC MF stimulation at −70 mV (solid lines) and 0 mV (dashed lines). **H**, Example of slow outward current evoked after optogenetic stimulation of NC pathway at 20 Hz. **I**, Summary of fast phasic current amplitudes recorded at −70 or 0 mV. **J**, Summary of charge transfer during slow outward currents evoked by NC MF stimulation. **K-M**, Same as F-G but for pontine MF stimulation. **N**, Example of slow outward current evoked after optogenetic stimulation of pontine MFs at 20 Hz. **O**, Same as I but for pontine MF stimulation. **P**, Same as J but for pontine MF stimulation.

EPSCs evoked from optogenetic excitation of the NC had an average amplitude of 80.6 ± 9.3 pA (Fig. 4B, F; sem; n = 7) and 10-90% rise and 37% decay times typical of AMPA type glutamate receptor-mediated currents in granule cells, averaging 0.25 ± 0.04 ms and 1.0 ± 0.2 ms respectively, similar to group 2 excitatory inputs previously described (Chabrol et al., 2015). Surprisingly, however, given the large spatial convergence of MFs and subsequent divergence of inhibition within the GCL fast phasic IPSCs recorded at 0-3 mV holding potentials were uncommon, observed in just 5/12 responsive GrCs and 5/96 total GrCs (Fig. 4C, G). Fast phasic IPSCs were distinct from EPSCs, with smaller amplitudes and slower kinetics, averaging 40.3 ± 5.5 pA, with 0.9 ± 0.1 ms 10-90% rise times and 2.7 ± 0.4 ms 37% decay times (Fig. 4I; n = 5; EPSC vs IPSC amplitude, p=0.007; rise time, p < 0.001; decay time, p = 0.001; unpaired t-tests). These measurements were statistically indistinguishable from spontaneous fast phasic inhibitory currents (amplitude, p = 0.1; rise time, p = 0.3; decay time, p = 0.2; unpaired t-tests), suggesting they originate from single GoCs rather than recruitment of multiple convergent GoCs. Moreover, as expected for feedforward inhibition, IPSC latencies and temporal variability were distinct from those of EPSCs (Fig. 4D), averaging 13.5 ± 0.9 ms vs 4.2 ± 0.3 ms for EPSCs, with latency jitter (SD of latency) averaging 3.6 ± 0.7 ms versus 0.29 ± 0.2 ms for EPSCs (Fig. 4E; p < 0.001; unpaired t-test). Thus, fast phasic inhibition observed in GrCs following NC optogenetic stimulation had the hallmark of feedforward inhibition mediated through GoC recruitment but was nevertheless uncommon.

Given the shared sparseness of direct excitatory and disynaptic phasic inhibitory inputs to GrCs following NC stimulation, we next examined the overlap of these inputs onto single GrCs. As expected for sparse synaptic contacts, GrCs with phasic responses to NC pathway stimulation showed either EPSCs (58.3%) or fast phasic IPSCs (41.6%) but not both in our recording set (Fig. 4I). When NC EPSCs were elicited at −70 mV, no fast phasic IPSC was detectable holding the cell at 0 mV; conversely, when IPSCs were evident at 0 mV, no EPSC was detectable at −70 mV.

Slow spillover mediated inhibition has been proposed as the primary form of inhibition within the GCL, so we next examined GrCs for slow inhibitory currents. Slow inhibitory currents were observed in all GrCs with phasic excitatory or inhibitory responses, with an average charge transfer of 124.5 ± 21 (pA·ms; IPSC area) (Fig. 4H, J). Thus, slow phasic inhibition dominates feedforward inhibitory processing from identified MF pathways, consistent with observations from electrical stimulation.

To contrast experiments studying low density of MFs from the NC pathway, we next examined GrC responses to a dense MF population originating in the pontine nuclei with optogenetics (Huang et al., 2013; Gilmer and Person, 2017). We recorded whole cell patch clamp recordings of GrCs in the vicinity of labeled MFs. Light stimulation evoked excitatory or fast phasic inhibitory currents in 21 GrCs (Fig. 4K, L, M; EPSCs, 55.6 ± 7.6 pA, with 0.4 ± 0.03 ms 10-90% rise times and 1.3 ± 0.11 ms 37% decay times, 2.8 ± 0.2 ms latency (n=18); IPSCs; 22.2 ± 1.2 pA, with 2.2 ± 0.4 ms 10-90% rise times and 2.3 ± 0.5 ms 37% decay times; 12.8 ± 2.1 ms latency (n=8)). The fraction of overlap of phasic EPSCs and fast phasic IPSCs was higher with pontine stimulation, as expected of a denser input: EPSCs without phasic IPSCs were evoked in 62% of GrCs and exclusively phasic IPSCs were evoked in 14% of recorded GrCs, constituting 76% of responsive cells. In the remaining 24% of responsive neurons, pontine MF stimulation evoked both EPSCs and fast phasic IPSCs (5/21) (Fig. 4O). Thus, density of MF afferents influences the overlap between phasic excitation and inhibition. As was observed for NC fibers, however, slow inhibitory currents dominated fast phasic inhibition, and were seen in all cells with phasic currents following pontine MF activation, with an average charge transfer of 88 ± 8.0 (pA ∙ ms) (Fig. 4N, P).

To summarize, fast phasic inhibition was probabilistically recruited onto GrCs depending on the approximate density of MF terminals but slow inhibitory currents were much more widespread, even when evoked by a sparse MF input.

### Limited role for fast phasic FFI in regulating GrC response timing

GoC-mediated feedforward inhibition has been proposed to shorten GrC responses to MF input via a temporal windowing effect, analogous to functions in neocortical circuitry (Pouille and Scanziani, 2001). We directly tested this idea in GrCs, combining optogenetic stimulation of pontine MFs and dynamic clamp, mimicking physiologically realistic feedforward fast and slow phasic inhibitory conductances. We recorded from GrCs in the presence of SR95531 (10 μM) to block endogenous GABA_a_ receptors. In whole cell current clamp mode, we measured GrC firing in response to 20 Hz optogenetic stimulation of pontine MFs, followed by delayed fast phasic IPSGs or slow phasic IPSGs produced via the dynamic clamp, mimicking GoC-mediated feedforward inhibition. GrCs were held at −60 to −75 mV by current injection to promote MF-driven firing. We used two conductances for both fast and slow phasic currents, with the lower of the two matching physiologically measured values but the higher providing insight into the upper bound of physiological levels of inhibition.

Low conductance fast phasic IPSGs (0.3 nS; rise time 2.15 ms, decay tau 2.29 ms), approximating a single fast phasic GoC input, were delayed by 12.8 ms relative to light in a 20 Hz light pulse 500 ms train to approximate feedforward inhibition. Across the population, this manipulation did not significantly change optogenetically driven firing compared to optogenetic stimulation alone (Fig. 5A, B; −3.9 ± 2.3 Hz from baseline, p > 0.1, paired t-test; n=8), although in one cell fast phasic IPSGs significantly reduced MF-driven firing responses (1/8 cell; −10.8 Hz from baseline, pontine 81.8 ± 2.0 sp/s, pontine + IPSG 71.0 ± 1.8 sp/s, unpaired t-test, n=10 pontine; n=5 pontine + IPSG trials; p=0.002). As a positive control, we next tested whether stronger phasic inhibition influenced firing rate. We increased the magnitude of the phasic stimulation to 1.2 nS, which is physiologically unlikely, since fast phasic contacts are rare and this assumes 4 onto a single cell, when only 60% of GrC dendrites receive direct contacts (Jakab and Hamori, 1988; Rossi and Hamann, 1998). As expected, this manipulation reduced response frequency (Fig. 5D, E; −11.9 ± 3.4 Hz from baseline, p < 0.05, paired t-test, n=8), with significant effects seen in 4/8 cells (p<0.05, unpaired t-test, n=10 pontine; n=5 pontine + IPSG trials in each cells). The temporal windowing hypothesis predicts that FFI shortens the duration of spiking in response to excitation. We therefore next measured the duration of firing in response to optogenetic activation of MFs with and without phasic IPSGs. Neither small nor large conductance fast phasic IPSGs significantly reduced the duration of firing across the population (Fig. 5C, F, low: pontine 377.5 ± 41.0 ms, pontine + IPSG 403.3 ± 52.7 ms, p=0.4, paired t-test, n=8, high: pontine 409.6 ± 25.2 ms, pontine + IPSG 333.4 ± 45.0 ms, p=0.07, paired t-test, n=8). This indicates that feedforward fast phasic inhibition of GrCs plays little role in shortening GrC responses to MF inputs.

**Figure 5.**
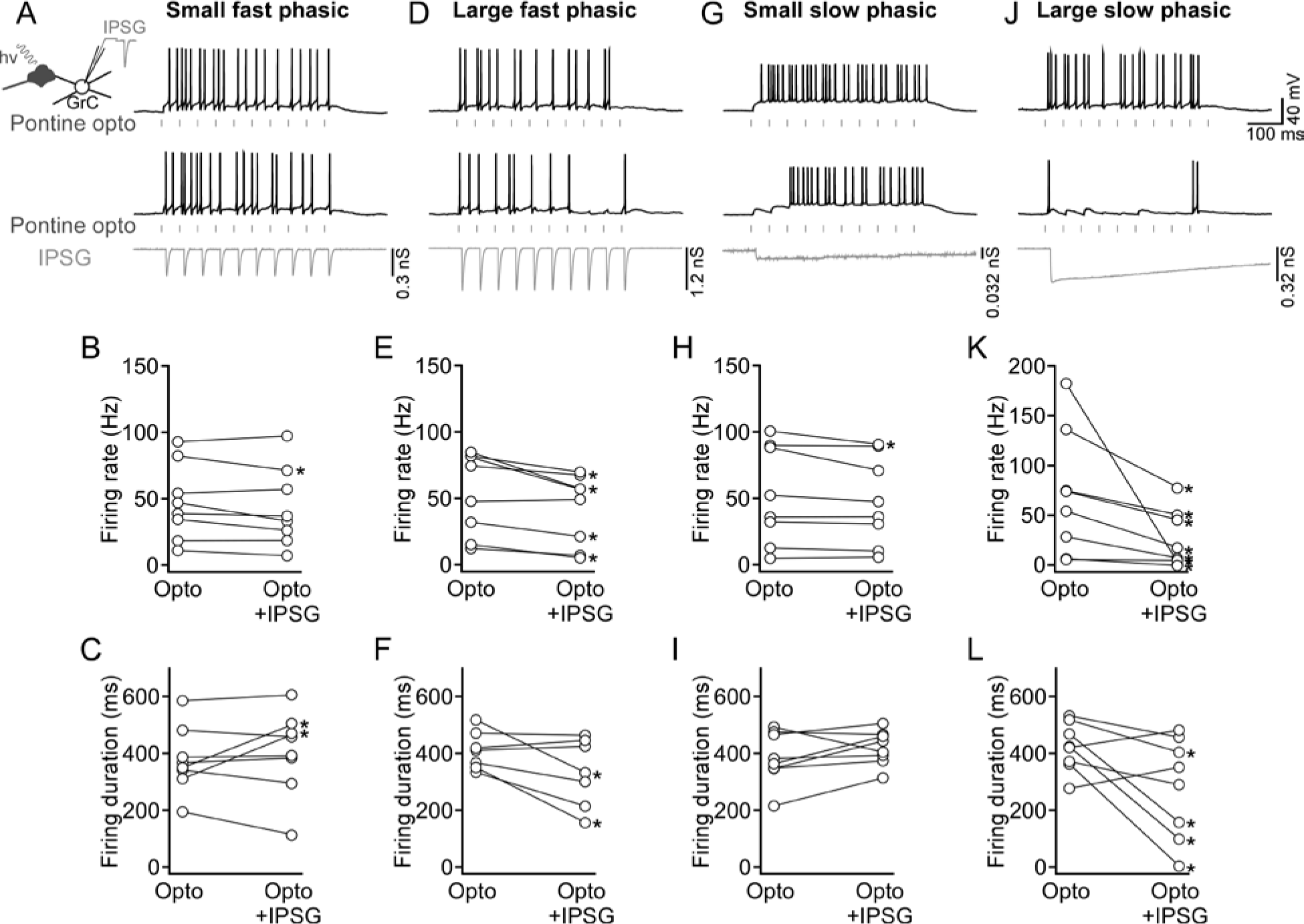
Predominant role for slow feedforward inhibition in regulating GrC synaptic responses. **A**, *Left*, Schematic of recording configuration where dynamic clamp and optogenetic MF stimulation were combined to examine the role of feedforward inhibition (A). *Right*, Representative traces showing GrC responses to 20 Hz optogenetic stimulation of pontine MFs without (upper trace) and with (lower trace) dynamic clamp physiological fast phasic IPSGs (lower). **B**, Summary of MF-evoked firing rate changes with and without small fast phasic IPSGs. **C**, Summary of MF-evoked firing durations with and without small fast phasic IPSGs. **D-F**, Same as A-C except with large fast phasic IPSGs. **G-I** Same as A-C except with small, slow phasic IPSGs. **J-L**, Same as A-C except with large, slow phasic IPSGs. Significant changes in firing rate were observed in 7/8 GrCs (p<0.05, paired t-tests). Significant changes in firing duration were observed in 4/8 GrCs (p<0.05, paired t-test).

In previous dynamic clamp studies, slow spillover-like inhibitory currents attenuated GrC firing more effectively than fast phasic IPSCs (Crowley et al., 2009). We extended these experiments to explicitly test a role for feedforward inhibition, such that the slow current was delayed relative to excitation and used conductances mimicking current from a single GoC or higher conductances that reflect summated slow phasic inhibition. We first mimicked slow phasic inhibition from a single GoC, injecting IPSGs (0.032 nS) following the first light stimulation, again delayed by 12.8 ms. This experimental condition did not reduce MF driven firing rates (Fig. 5G, H, −4.4 ± 2.2 Hz from baseline, p >0.05, paired t-test, n=8). Because multiple GoCs converge in a glomerulus we next tested higher conductance IPSGs that reflect summated slow phasic inhibition. We therefore increased conductance 10 fold capturing summation. As expected, larger slow phasic IPSGs (0.32 nS) strongly attenuated rates in most GrCs (Fig. 5J, K, 7/8 cells, p<0.05, unpaired t-test, n=10 pontine; n=5 pontine + IPSG trials in each cells); however as was seen with fast phasic IPSGs, neither low nor high conductance slow IPSGs had significant effects on firing duration (Fig. 5 I, L, pontine, 391.7 ± 32.5 ms, pontine + 0.032 nS IPSG, 425.9 ± 21.6 ms, p=0.18, paired t-test, n=8; pontine + 0.32 nS IPSG, 278.9 ± 62.6 ms, p=0.053, paired t-test, n=8). These data illustrate a preferential role for convergent GoC activity on GrC firing rates but not overall firing duration in response to excitatory inputs, and do not support a temporal windowing role for feedforward inhibition by GoCs.

### GoC recruitment scales with MF input density

The dynamic clamp experiments indicate that fast phasic feedforward inhibition from GoCs plays only a minor role in temporally structuring GrC responses to MF input and support the alternative view that slow, convergent inhibitory events preferentially regulate the threshold of GrCs (Brickley et al., 1996; Wall and Usowicz, 1997; Rossi and Hamann, 1998; Hamann et al., 2002; Mitchell and Silver, 2003). Threshold regulation was a major role for GoCs in theoretical studies proposing a role for pattern discrimination by GrCs, and predict that GoC recruitment would scale with input density (Marr, 1969; Pellionisz and Szentagothai, 1973; Mapelli et al., 2009; Honda and Ito, 2017). Therefore, identifying rules of GoC recruitment is essential to differentiate between diverse models of GrC information processing.

To better understand the rules of GoC recruitment by identified MF inputs, we next recorded from GoCs directly, measuring evoked firing and synaptic currents following optogenetic stimulation of MFs originating in the cerebellar or pontine nuclei (Fig. 6A). We used GlyT2-eGFP mice to identify GoCs under fluorescence, noting that this excludes a small subpopulation of GoCs from our dataset (Simat et al., 2007). We first tested the efficacy of EPSPs from both NC and pontine sources to drive firing in GoCs in current clamp mode during trains of stimuli delivered at rates between 20-60 Hz (Fig. 6C). We performed linear regression analysis on firing rate changes as a function of stimulation frequency. Consistent with previous observations from electrical stimulation (Kanichay and Silver, 2008), MF stimulation rates were poor predictors of firing rate changes across the population (R^2^ = 0.03, p= 0.03, F= 4.6, df= 176, n=178 from 36 cells pooled).

**Figure 6.**
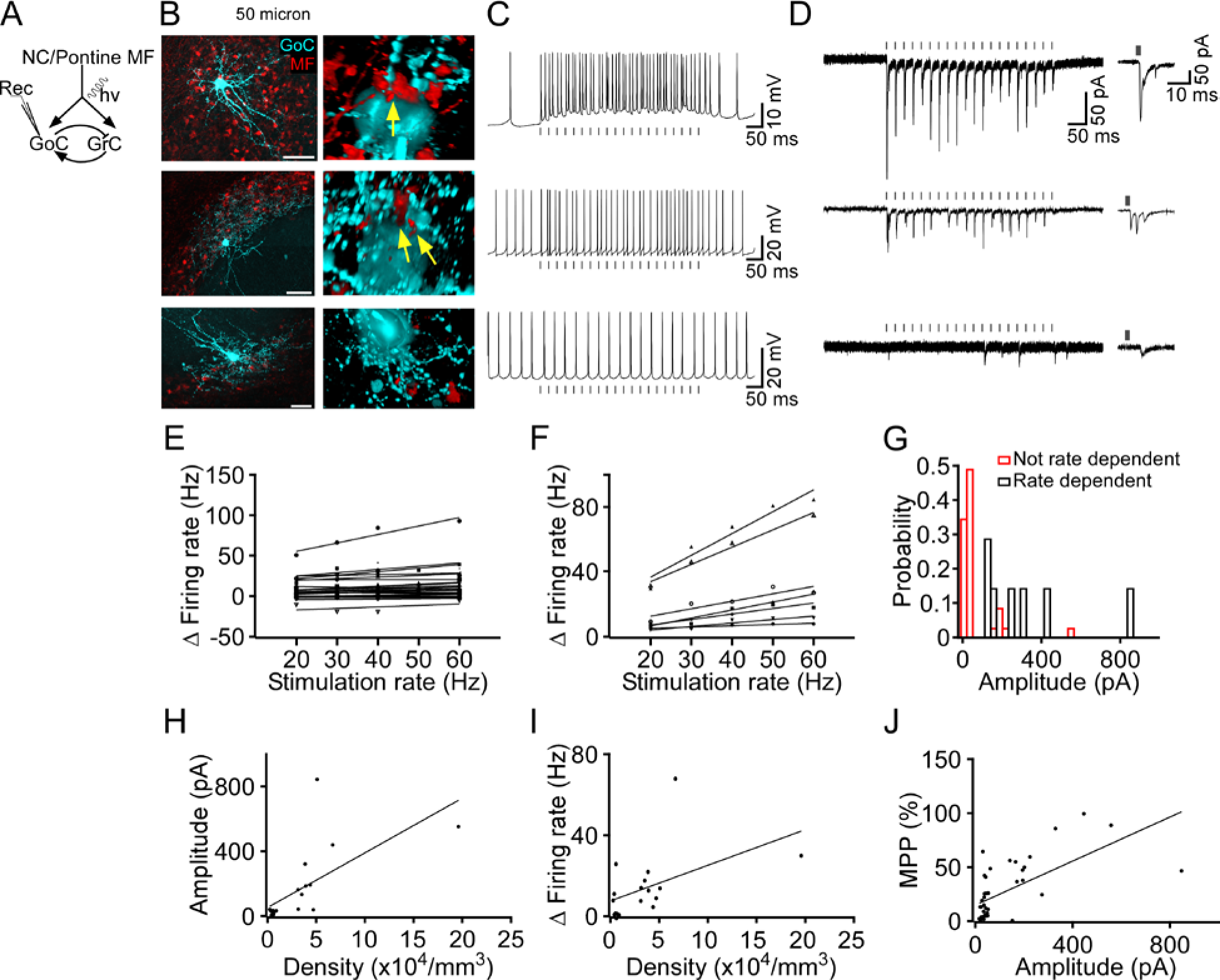
GoC recruitment follows MF population activity levels. **A**, Schematic diagram of recording configuration. ChR2-expressing NC or Pontine MFs were stimulated during GoC recordings. **B**, Biocytin filled GoCs (cyan) recovered after recordings show proximity of RFP-expressing MFs. Scale: 50 μm. Right, Putative synaptic contacts (arrows) or absence of contacts between recorded GoC and MF, associated with physiological traces in C and D shown in transparency rendering mode. **C**, Representative examples of GoC evoked firing in response to optogenetic stimulation of pontine MFs at 40 Hz. Responses varied between cells (top-bottom). **D**, Representative traces showing diversity of evoked EPSCs following optogenetic stimulation of pontine MFs at 40 Hz. Traces in D are matched with current clamp responses in C. **E and F**, Relationship of GoC firing rate change to stimulation rate. Non-significant relationships are shown in E, Significant relationships are shown in F. **G**, Probability distribution of initial EPSC amplitudes measured in GoCs showing significant (black; data from F) or non-significant (red; data from E) input-output relationships. **H**, Relationship of EPSC amplitude to ChR2-expressing MF density. (R^2^= 0.44, p = 0.001, F=14.3, df=18, n=20). **I**, Relationship of GoC firing rate change to 40 Hz stimulation as a function of ChR2-expressing MF density. (R^2^ = 0.23; p=0.04; F= 5.0; df= 16; n= 18). **J**, Multi-EPSC-peak probability plotted as a function of initial EPSC amplitude. (R^2^=0.41, p<0.0001, F=27.9, DF=40, n=42).

This filtering property has been previously ascribed to a large afterhyperpolarization from EPSP-driven spikes (Kanichay and Silver, 2008). Nevertheless, it raises the question of how GrC population activity could remain constant if GoC recruitment is strongly filtered, prompting us to look closer at the relationship between MF activity and GoCs. Indeed, when we examined the relationship of firing rate with stimulation rate in individual neurons, we noted that some were strongly correlated, with significant within-cell correlation (Fig. 6F; p<0.04, n=7) while many cells were not (Fig. 6E; p>0.05, n=28). We next examined whether properties of the EPSCs differed between these groups and found a strong bias toward stronger EPSCs supporting linear input-output relations (Fig. 6G; 332.1 ± 41.5 pA, n=7). Weaker EPSCs, by contrast, did not modulate GoC spike rates with increasing stimulus rates (Fig. 6G; 71.5 ± 8.1 pA, n=35).

These findings indicate that EPSC amplitude determines the input filtering properties of GoCs. To identify factors that influence the amplitude of EPSCs onto GoCs, we took advantage of the fact that optogenetic probes are coupled with a fluorophore, thus we could directly measure the density of activated inputs. This allowed us test the prediction that MF convergence onto GoCs from multiple inputs would summate to regulate EPSC amplitude in GoCs. Consistent with high levels of convergence of MFs onto GoCs, there was a strong linear relationship between MF density and EPSC amplitude, with higher density MFs eliciting larger EPSCs (Fig. 6H; R^2^ = 0.44, p=0.001, F=14.3, df=18, n=20). Despite the sparseness of NC MFs, which constitute just 1-10% of local MFs (Gilmer and Person, 2017), optogenetic activation of the pathway at 40 Hz elicited EPSCs in 35% of GoCs recorded in the vicinity of labeled MFs (34/96). Denser MF populations from the pontine nuclei elicited responses in 87% of nearby GoCs (14/16). Not surprisingly, given the large difference in density between pontine and NC inputs pontine inputs elicited significantly larger EPSCs (p<0.001, unpaired t-test, −259.5±26.8 pA, n=70 from 14 cells pooled, −40.7±3.1 pA, n=131 from 28 cells pooled, respectively). These observations suggest that GoCs integrate many inputs from diverse sources and are responsive to the level of input population activity as expected for active thresholding.

Having identified a relationship between MF input density and GoC EPSC amplitude and linearity of firing responses, we next reasoned that GoC firing responses might be sensitive to the overall level of MF activity levels, i.e. the density of active inputs. We tested this idea by analyzing the density of MFs labeled within the vicinity of the recorded GoC, and relating measurements to the magnitude of firing rate changes observed with 40 Hz stimulation. In keeping with this reasoning, we found a positive significant correlation between active input density and the magnitude of firing rate changes during stimulation (Fig. 6I; R^2^ = 0.23; p=0.04; F= 5.0; df= 16; n= 18).

As described in previous studies (Kanichay and Silver, 2008; Hull and Regehr, 2012; Cesana et al., 2013; Gao et al., 2016), there was considerable diversity in GoC EPSC properties, which differed by amplitude, latency and whether they were singular or included disynaptic feedback via parallel fiber input from GrCs. In general, EPSCs fell into three broad classes, which included (1) short latency; (2) mixed short- and long-latency producing multiple peaks per stimulus; (3) and long-latency (Fig. 6D). Consistent with the view that these response classes reflect mono and di-synpatic input, biocytin fills of recorded GoCs revealed ChR2-RFP expressing MF inputs adjacent to somata and/or basal dendrites in the case of short-latency (<3.6 ms) responders (9/11 short latency responders were recovered) but not in the case of long-latency responders (3/3 > 3.8 ms; Fig. 6B). Furthermore, bath application of the NMDA receptor antagonist CPP reduced the likelihood of multipeaked EPSC being elicited from 37.9 ± 14.0% to 5.8 ± 1.0 % (n=3), supporting the view that late synaptic responses were the result of disynaptic recruitment of GrCs (Cesana et al., 2013).

Because GoC multipeaked EPSCs are a readout of GrC recruitment which is in turn regulated by GoC inhibitory feedback (Cesana et al., 2013), we reasoned that we could use the relationship between MF input strength and multipeaked EPSC probability to test whether GoCs dynamically modulate MF-GrC gain as predicted in numerous models (Mitchell and Silver, 2003). GoC EPSC multipeak probability was linearly correlated with the initial EPSC amplitude (Fig. 6J; R^2^=0.41, p<0.0001, F=27.9, DF=40, n=42), indicating that stronger input recruited more GrCs. Interestingly, however, there was no detectable change in multipeak probability or number of peaks over the course of moderate frequency (40 Hz) stimulus trains (p = 0.68, p = 0.65, paired t-test of early vs late train multipeak probability or early vs late train number of peaks per stimulus). This stability of multipeak probability was evident regardless of whether multipeak probability was high or low at the beginning of the stimulus train. Coupled with the observation that GABA_A_ receptor blockade strongly enhances multipeak probability (Cesana et al., 2013), these findings indicate that feedback inhibition within the layer stabilizes GrC excitability for a given input excitation level.

## Discussion

This study used a combination of quantitative morphometry, optogenetics and dynamic clamp to test theoretical predictions about GoCs as global regulators of GrC excitability. Theory has long posited a role for GoCs in regulating GrC population activity (Eccles et al., 1967; Marr, 1969), proposing that they respond dynamically to varying input levels to modulate all local GrC thresholds, but data testing key assumptions of this view have been lacking. Here we confirm theoretical predictions that GoC axons are ideally suited to globally inhibit neighboring GrCs; that spillover mediated feedforward inhibition preferentially alters GrC thresholds; and that convergent afferents are essential to recruit GoC inhibition in a manner consistent with a global ‘listening’ mechanism suited for normalizing activity in the GCL. These data indicate that the large multimodal integrative capacity of the GoC, combined with physiological integrative rules of GrCs set up a MF activity level-detector to regulate inhibition levels within the GCL. These data disfavor a role for GoCs in temporally sharpening GrC responses to MF input, simplifying the computational role of GoCs in the layer.

GrCs have been proposed to perform pattern separation by sparsening information conveyed by MF inputs (Cayco-Gajic et al., 2017). Inhibition from widefield interneurons has been proposed in both mammalian cerebellum and other systems to perform these computations (Pouille et al., 2009), and assume that inhibitory interneurons contact all or nearly all neurons in the field to effectively regulate population activity (Marr, 1969; Albus, 1971; Billings et al., 2014; Duguid et al., 2015; Cayco-Gajic et al., 2017). Similar roles have been proposed for cerebellar GoCs but although their morphology has long been appreciated as complex and suitable for widespread inhibition, quantitative analysis of axonal density has not been performed, leading to conflicting speculation about whether inhibition they produce is spatially structured. Our morphometry shows that GoCs are suited to contact every nearby GrC. This is important in light of potential alterative computations that could distribute inhibition to subsets of neighboring GrCs to decorrelate or temporally sculpt responses. Our findings show that rather, cerebellar GoCs favor a blanket inhibitory process within the region.

Our quantitative morphometry of reconstructed GoCs allowed us to evaluate how a population of GoCs might contribute to spatial distribution of inhibition. While Golgi cell axonal fields vary in their orientation and extent (Simpson et al., 2005; Holtzman et al., 2006; Barmack and Yakhnitsa, 2008) they share dense axonal arborization across cells. While partial reconstruction of cerebellar GoCs have been published, they have most often been following labeling in brain slices which necessarily exclude processes leaving or re-entering the slice (Kanichay and Silver, 2008; Vervaeke et al., 2010; Vervaeke et al., 2012; Szoboszlay et al., 2016; Valera et al., 2016) or have been incomplete reconstructions of in vivo cell labeling (Simpson et al., 2005; Barmack and Yakhnitsa, 2008).

The comprehensive reconstructions performed here offer insight into a GoC structural motif, where boutons from individual cells remain tightly clustered even though the overall density of boutons falls off with distance from the soma. Measurements of GlyT2-positive GoC bouton density in similar preparation allowed direct comparison of individual GoCs within the broader population, and suggest that within 50 μm of a GoC soma, single local GoCs dominate the inhibitory axonal population, with minor contributions from approximately 5 overlapping GoCs. This may be an underestimate owing to a remaining small population of neurogranin positive GlyT2 negative GoCs (Simat et al., 2007), but is in line with physiological estimates (Rossi and Hamann, 1998). The density of local boutons suggests multiple innervation of every glomerulus by individual GoCs in the field. It is important to note that average density measurements do a poorer job of resolving the overlap of neighboring GoCs, since they do not take into account local clustering. Based on estimates of glomerular density (Palkovits et al., 1971; Billings et al., 2014), our measurements would suggest that on average an individual GoC makes between 0.5 and 1.5 boutons per glomerulus within its axonal field, with numbers fluctuating depending on local clustering. Jakab and Hamori (1988) measured 12-28 GoC boutons per glomerulus in cat, which roughly aligns to the 20-fold difference in single vs population bouton density in our measurements, which overestimates GoC convergence.

The conclusions drawn from the reconstruction data indicate that the GoC is in a position to ‘globally’ inhibit GrCs within range of its axonal arbors, as seen in thresholding motifs (Marr, 1969; Albus, 1971). However, examination of GrCs that were excited by to optogenetic activation of MF input revealed surprisingly sparse fast phasic inhibition, even when MF inputs were dense (Fig. 4). This seeming inconsistency was resolved by analyzing GrCs for spillover-like slow inhibitory currents. This form of inhibition, which differs in kinetics and amplitude from direct phasic inhibition was always observed in GrCs activated by MFs. Slow spill-over mediated IPSCs have been extensively studied in GrCs (Rossi and Hamann, 1998; Mitchell and Silver, 2000; Rossi et al., 2003; Duguid et al., 2012; Duguid et al., 2015) and shown to dominate inhibitory processing in GrCs. The difference in likelihood of fast phasic and slow spillover currents, coupled with the extremely high density of GoC boutons suggested that fast phasic inhibitory events occur probabilistically due to the chance spatial proximity of GrCs dendrites relative to GoC axonal boutons. This view may be consistent with the observation that ‘tonic inhibition’ is present in the GCL. This form of inhibition, which lacks resolvable inhibitory current kinetic modulation, may be at the end of a continuum of inhibitory currents where the distance from GrC dendrite to GoC boutons releasing GABA dictates kinetics.

Nevertheless, the striking differences between these IPSCs prompted further investigation into the relative physiological roles of these forms of inhibition. Previous studies have shown using dynamic clamp that slow inhibition strongly inhibits GrCs (Crowley et al., 2009; Solinas et al., 2010; Duguid et al., 2015). We extended these studies by combining physiological levels of inhibition through a dynamic clamp with optogenetic activation of pontine MFs to directly test the temporal windowing hypothesis of phasic inhibition (Kalmbach et al., 2011). We specifically tested if phasic inhibition or slow spillover inhibitory events, offset in time from EPSPs, as would occur with feedforward inhibition, modulated the duration of spiking in GrCs. Surprisingly, we found little consistent effect of fast phasic inhibition on any parameter of GrC spiking - with no consistent effects on spike rate or duration of firing. By contrast, slow spillover like conductances, offset in time relative to opto-EPSPs did reduce rate, but effects on spiking duration were inconsistent.

It was surprising to us to find such a limited effect of phasic inhibition on the timing of spiking in GrCs, particularly since regulating the duration of MF mediated bursting would seem to be a straight forward mechanism of sparsening GrC recruitment or diversifying spike timing. However, recent work has proposed an alternative mechanism based on diversity in short-term plasticity of MFs (Chabrol et al., 2015), thus GoC mediated FFI need not be in place to accomplish temporal diversification of GrC firing.

Another source of complexity in relating MF activity to recruitment of feedforward inhibition is the fact that EPSP-spike coupling in GoCs is heavily temporally filtered (Vervaeke et al., 2010; Kanichay and Silver, 2008). Our study corroborated reports from several groups finding that GoCs do not necessarily follow increasing MF stimulation rates with high fidelity. Because our study involved optogenetic recruitment of MFs labeled with a fluorophore, we could relate GoC activation in our physiology recordings to the density and distribution of MFs activated by light. We uncovered a linear relationship between active MF density and EPSC amplitudes in GoCs elicited by light stimulation, indicative of convergence onto GoCs. Furthermore, density also related to the amplitude of firing elicited at a fixed stimulus rate. In contrast to highly filtered EPSP-GoC firing rate relationships seen across the population and with electrical stimulation, firing rates of GoCs that received the strongest EPSCs from optogenetic stimuli (i.e. those with the highest density of MFs) linearly increased with stimulation rates. These findings relate the density of MF activity to the recruitment of GoCs in a rate dependent manner, and suggest inhibitory mechanisms within the GCL maintain a large dynamic range by avoiding saturation from sparse MF activation via nonlinear recruitment. The statistical structure of feedback inhibition is important for models of GrC inhibitory regulation, where, depending on the physical structure of the cell in relation to its inputs, could form one of several circuit motifs that mediate alternative computations (Braganza and Beck, 2018). GoCs resemble widefield interneurons in insect olfactory system that sample input density and widely inhibit target populations. Interestingly, the insect mushroom bodies possess other circuit motifs similar shared in cerebellum, such as convergent, locally random inputs to neurons with sparse dendrites, resembling cerebellar GrCs (Litwin-Kumar et al., 2017). That the circuits share numerous structural motifs underlines the potential commonality to their computations and suggests convergent evolution.

In conclusion this study relates GoC morphology and unique synaptic physiology onto GrCs to local circuit computations that function to sparsen GrC activity relative to MF inputs by reducing excitability. Our data reveal a strikingly dense inhibitory field of local GoCs, analogous to the widefield inhibitory interneurons in insect mushroom bodies (Papadopoulou et al., 2011) and show that individual GoCs integrate many MF inputs to regulate their output, driving activity that scales with input. Thus, the GoC occupies a key multimodal integrative niche within the layer that, compared to the extremely limited extent of the GrC dendrite, allows for a broader integration of multimodal signals to regulate GrC population activity.

## Conflict of Interest

The authors declare no competing financial interests.

## Acknowledgements

We thank Ms Samantha Lewis for expert technical support during the project. This work was supported by the Japan Society for The Promotion of Science (JSPS) Overseas Research Fellowship and The Uehara Memorial Foundation research fellowship to S.T.; NS084996; a Kingenstein Foundation fellowship; and a Boettcher foundation Webb-Waring biomedical research award to A.L.P. Imaging experiments were performed in the University of Colorado Anschutz Medical Campus Advance Light Microscopy Core supported in part by Rocky Mountain Neurological Disorders Core Grant Number P30NS048154 and by NiH/NCATS Colorado CTSI Grant Number UL1 TR001082. Engineering support was provided by the Optogenetics and Neural Engineering Core at the University of Colorado Anschutz Medical Campus, funded in part by the National Institute of Neurological Disorders and Stroke of the National Institutes of Health under award number P30NS048154.

## References

Albus JS (1971) A theory of cerebellar function. Mathematical Biosciences 10:25–61.

Ankri L, Husson Z, Pietrajtis K, Proville R, Lena C, Yarom Y, Dieudonne S, Uusisaari MY (2015) A novel inhibitory nucleo-cortical circuit controls cerebellar Golgi cell activity. Elife 4.

Barmack NH, Yakhnitsa V (2008) Functions of interneurons in mouse cerebellum. J Neurosci 28:1140–1152.

Billings G, Piasini E, Lorincz A, Nusser Z, Silver RA (2014) Network structure within the cerebellar input layer enables lossless sparse encoding. Neuron 83:960–974.

Braganza O, Beck H (2018) The Circuit Motif as a Conceptual Tool for Multilevel Neuroscience. Trends Neurosci 41:128–136.

Brickley SG, Cull-Candy SG, Farrant M (1996) Development of a tonic form of synaptic inhibition in rat cerebellar granule cells resulting from persistent activation of GABAA receptors. J Physiol 497 (Pt 3):753–759.

Cayco-Gajic NA, Clopath C, Silver RA (2017) Sparse synaptic connectivity is required for decorrelation and pattern separation in feedforward networks. Nat Commun 8:1116.

Cesana E, Pietrajtis K, Bidoret C, Isope P, D’Angelo E, Dieudonne S, Forti L (2013) Granule cell ascending axon excitatory synapses onto Golgi cells implement a potent feedback circuit in the cerebellar granular layer. J Neurosci 33:12430–12446.

Chabrol FP, Arenz A, Wiechert MT, Margrie TW, DiGregorio DA (2015) Synaptic diversity enables temporal coding of coincident multisensory inputs in single neurons. Nat Neurosci 18:718–727.

Crowley JJ, Fioravante D, Regehr WG (2009) Dynamics of fast and slow inhibition from cerebellar golgi cells allow flexible control of synaptic integration. Neuron 63:843–853.

D’Angelo E, De Zeeuw CI (2009) Timing and plasticity in the cerebellum: focus on the granular layer. Trends Neurosci 32:30–40.

D’Angelo E, Solinas S, Mapelli J, Gandolfi D, Mapelli L, Prestori F (2013) The cerebellar Golgi cell and spatiotemporal organization of granular layer activity. Front Neural Circuits 7:93.

Desai NS, Gray R, Johnston D (2017) A Dynamic Clamp on Every Rig. eNeuro 4.

Duguid I, Branco T, London M, Chadderton P, Hausser M (2012) Tonic inhibition enhances fidelity of sensory information transmission in the cerebellar cortex. J Neurosci 32:11132–11143.

Duguid I, Branco T, Chadderton P, Arlt C, Powell K, Hausser M (2015) Control of cerebellar granule cell output by sensory-evoked Golgi cell inhibition. Proc Natl Acad Sci U S A 112:13099–13104.

Eccles JC, Ito M, Szentagothai J (1967) The Cerebellum as a Neuronal Mechine. Berlin: Springer-Verlag.

Farrant M, Nusser Z (2005) Variations on an inhibitory theme: phasic and tonic activation of GABA(A) receptors. Nat Rev Neurosci 6:215–229.

Gao Z, Proietti-Onori M, Lin Z, Ten Brinke MM, Boele HJ, Potters JW, Ruigrok TJ, Hoebeek FE, De Zeeuw CI (2016) Excitatory Cerebellar Nucleocortical Circuit Provides Internal Amplification during Associative Conditioning. Neuron 89:645–657.

Gilmer JI, Person AL (2017) Morphological Constraints on Cerebellar Granule Cell Combinatorial Diversity. J Neurosci 37:12153–12166.

Hamann M, Rossi DJ, Attwell D (2002) Tonic and spillover inhibition of granule cells control information flow through cerebellar cortex. Neuron 33:625–633.

Holtzman T, Rajapaksa T, Mostofi A, Edgley SA (2006) Different responses of rat cerebellar Purkinje cells and Golgi cells evoked by widespread convergent sensory inputs. J Physiol 574:491–507.

Honda T, Ito M (2017) Development from Marr’s theory of the cerebellum. In: Computational theories and their implementation in the brain: the legacy of David Marr (Vaina LM and Passingham RE, ed), First edition. Edition. Oxford: Oxford University Press.

Huang CC, Sugino K, Shima Y, Guo C, Bai S, Mensh BD, Nelson SB, Hantman AW (2013) Convergence of pontine and proprioceptive streams onto multimodal cerebellar granule cells. Elife 2:e00400.

Hull C, Regehr WG (2012) Identification of an inhibitory circuit that regulates cerebellar Golgi cell activity. Neuron 73:149–158.

Ishikawa T, Shimuta M, Hausser M (2015) Multimodal sensory integration in single cerebellar granule cells in vivo. Elife 4.

Jakab RL, Hamori J (1988) Quantitative morphology and synaptology of cerebellar glomeruli in the rat. Anat Embryol (Berl) 179:81–88.

Kalmbach BE, Voicu H, Ohyama T, Mauk MD (2011) A subtraction mechanism of temporal coding in cerebellar cortex. J Neurosci 31:2025–2034.

Kanichay RT, Silver RA (2008) Synaptic and cellular properties of the feedforward inhibitory circuit within the input layer of the cerebellar cortex. J Neurosci 28:8955–8967.

Litwin-Kumar A, Harris KD, Axel R, Sompolinsky H, Abbott LF (2017) Optimal Degrees of Synaptic Connectivity. Neuron 93:1153–1164 e1157.

Mapelli L, Rossi P, Nieus T, D’Angelo E (2009) Tonic activation of GABAB receptors reduces release probability at inhibitory connections in the cerebellar glomerulus. J Neurophysiol 101:3089–3099.

Marr D (1969) A theory of cerebellar cortex. J Physiol 202:437–470.

Medina JF, Garcia KS, Nores WL, Taylor NM, Mauk MD (2000) Timing mechanisms in the cerebellum: testing predictions of a large-scale computer simulation. J Neurosci 20:5516–5525.

Mitchell SJ, Silver RA (2000) Glutamate spillover suppresses inhibition by activating presynaptic mGluRs. Nature 404:498–502.

Mitchell SJ, Silver RA (2003) Shunting inhibition modulates neuronal gain during synaptic excitation. Neuron 38:433–445.

Nieus TR, Mapelli L, D’Angelo E (2014) Regulation of output spike patterns by phasic inhibition in cerebellar granule cells. Front Cell Neurosci 8:246.

Palay SL, Chan-Palay V (1974) Cerebellar Cortex Cytology and Organication.

Palay SL, Chan-Palay V (1976) A guide to the synaptic analysis of the neuropil. Cold Spring Harb Symp Quant Biol 40:1–16.

Palkovits M, Magyar P, Szentagothai J (1971) Quantitative histological analysis of the cerebellar cortex in the cat. II. Cell numbers and densities in the granular layer. Brain Res 32:15–30.

Papadopoulou M, Cassenaer S, Nowotny T, Laurent G (2011) Normalization for sparse encoding of odors by a wide-field interneuron. Science 332:721–725.

Pellionisz A, Szentagothai J (1973) Dynamic single unit simulation of a realistic cerebellar network model. Brain Res 49:83–99.

Pouille F, Scanziani M (2001) Enforcement of temporal fidelity in pyramidal cells by somatic feed-forward inhibition. Science 293:1159–1163.

Pouille F, Marin-Burgin A, Adesnik H, Atallah BV, Scanziani M (2009) Input normalization by global feedforward inhibition expands cortical dynamic range. Nat Neurosci 12:1577–1585.

Rossi DJ, Hamann M (1998) Spillover-mediated transmission at inhibitory synapses promoted by high affinity alpha6 subunit GABA(A) receptors and glomerular geometry. Neuron 20:783–795.

Rossi DJ, Hamann M, Attwell D (2003) Multiple modes of GABAergic inhibition of rat cerebellar granule cells. J Physiol 548:97–110.

Rudolph S, Hull C, Regehr WG (2015) Active Dendrites and Differential Distribution of Calcium Channels Enable Functional Compartmentalization of Golgi Cells. J Neurosci 35:15492–15504.

Simat M, Parpan F, Fritschy JM (2007) Heterogeneity of glycinergic and gabaergic interneurons in the granule cell layer of mouse cerebellum. J Comp Neurol 500:71–83.

Simpson JI, Hulscher HC, Sabel-Goedknegt E, Ruigrok TJ (2005) Between in and out: linking morphology and physiology of cerebellar cortical interneurons. Prog Brain Res 148:329–340.

Solinas S, Nieus T, D’Angelo E (2010) A realistic large-scale model of the cerebellum granular layer predicts circuit spatio-temporal filtering properties. Front Cell Neurosci 4:12.

Szoboszlay M, Lorincz A, Lanore F, Vervaeke K, Silver RA, Nusser Z (2016) Functional Properties of Dendritic Gap Junctions in Cerebellar Golgi Cells. Neuron 90:1043–1056.

Valera AM, Binda F, Pawlowski SA, Dupont JL, Casella JF, Rothstein JD, Poulain B, Isope P (2015) Stereotyped spatial patterns of functional synaptic connectivity in the cerebellar cortex. Elife 5.

Vervaeke K, Lorincz A, Nusser Z, Silver RA (2012) Gap junctions compensate for sublinear dendritic integration in an inhibitory network. Science 335:1624–1628.

Vervaeke K, Lorincz A, Gleeson P, Farinella M, Nusser Z, Silver RA (2010) Rapid desynchronization of an electrically coupled interneuron network with sparse excitatory synaptic input. Neuron 67:435–451.

Wall MJ, Usowicz MM (1997) Development of action potential-dependent and independent spontaneous GABAA receptor-mediated currents in granule cells of postnatal rat cerebellum. Eur J Neurosci 9:533–548.

